# Non-intrusive optical measurement of egg geometry and volume for macro- and microscopic ovoids

**DOI:** 10.1101/2025.11.12.688065

**Authors:** Antonio Moreno-Rodenas, Sophie Le Hesran

**Author notes:** Correspondence should be addressed to S.L.H or A.M-R (;).

## Abstract

Egg size is a determining trait in the survival and development of bird and arthropod offspring. Ecologists studying these animal groups need reliable and practical methods to measure egg geometries. Despite the existence of numerous egg measurement techniques, the current scientific literature lacks a non-intrusive method allowing for field measurements without disrupting the measured individuals. Moreover, there has been so far no systematic comparison between the different available methodologies, making it difficult to assess their strengths and weaknesses. This study proposes a new and non-intrusive method for retrieving ovoid-shaped egg geometries, consisting in optimizing a parametric virtual model from N camera views (N-Views method). We tested the N-Views method on chicken (*Gallus gallus domesticus*) and quail (*Coturnix coturnix*) eggs, comparing its accuracy with 2 other existing optical measurement methods (Photogrammetry and Silhouette). Across all tested methods, N-Views provided the best estimate of egg volume when compared with the reference manual measurements (Archimedean buoyancy force), resulting in a MAE of 125 mm^3^ and 1.03% and 0.28% relative error for *Coturnix* and *Gallus* eggs, respectively. We further quantified the sensitivity of the N-Views and Silhouette methods to error sources arising from perspective and egg-camera alignment, highlighting the characteristics needed in optical measurement protocols for the estimation of egg geometries. Finally, we discuss the transferability of the N-Views method to the size estimation of microscopic arthropod eggs. This novel approach opens a new door for the non-intrusive exploration of relationships between egg size and life history traits in avian and arthropod species.

## Introduction

Reproductive success is an essential aspect in the life history of animal species, as natural selection favours individuals that leave the greatest number of descendants in future generations (Sinervo and Doughty, 1996; Giron and Casas, 2003). This reproductive success is largely based on offspring survival, a fitness component determining whether or not a juvenile will reach maturity and reproduce (Ronget et al., 2018). Understanding the ecological and biological factors affecting offspring survival is, therefore, a necessary step in the comprehension of population dynamics in animal species.

Offspring size is an especially interesting trait related to offspring survival and reproductive success, because it is simultaneously a maternal and offspring character (Fox and Czesak, 2000). In birds and arthropods, in particular, egg size is positively related to offspring survival (Giron and Casas, 2003; Bogdanova et al., 2006; Krist, 2011). Larger eggs provide a greater amount of nutrients and energy for growth, and progeny hatching from larger eggs can better withstand environmental stresses such as starvation or cold (Williams, 1994; Fox and Czesak, 2000; Ronget et al., 2018). On the maternal side, the age and physiological state of a female, as well as the environmental conditions she is exposed to, can affect the size of her eggs (Ronget et al., 2018). In birds, for example, young females have been found to lay smaller eggs than mature females (Giron and Casas, 2003). In arthropods, predatory mite females exposed to drought lay bigger eggs (Le Hesran et al., 2019).

Studying egg size requires reliable measurement methods. Traditional estimation of avian egg size has relied on manual techniques, using callipers to measure the maximum length and breadth, water displacement (Archimedes’ method) for volume estimation, or creating a tape envelope to estimate the surface area (Narushin, 2005; Zhang et al., 2016). However, these manual techniques are labor-intensive, intrusive (touching the egg, submerging it in fluids), unhygienic, subjective, error-prone and unsuitable for field conditions (Okinda et al., 2020). Moreover, they are restricted to relatively large eggs and cannot be transferred to microscopic eggs, common in arthropod species.

In recent years, several methods have been proposed to estimate egg geometries with the assistance of digital photography. A common approach (here referred to as the Silhouette method) estimates egg geometries using a 2D image of the egg contour (Soltani et al., 2015; Le Hesran et al., 2019; Shi et al., 2023). In the Silhouette method, a top-view image of the egg is taken, the contour of the egg is then delineated and length, volume, and surface area are estimated either parametrically or using the Pappus’ theorem for revolution solids. However, monocular 2D image-based analysis requires assumptions on the egg positioning (i.e., the main axis is aligned orthogonally with the image plane) and generally neglects camera projective errors. Moreover, aligning the egg with the camera plane is intrusive and impractical with microscopic eggs.

Other approaches have been proposed that acquire 3D point cloud estimates of the eggs to reconstruct their geometry, such as the use of photogrammetry (Zhang et al., 2016). Similarly, Chan et al., (2018) and Okinda et al., (2020) made direct distance measurements of the egg surface using infrared markers and light-ranging. These methods rely on multi-view approaches, but generally still require the egg to be manipulated and have certain properties (e.g. surface reflectivity or presence of features). Recently, Xiao et al., (2025) have also proposed a multi-view reconstruction approach for duck eggs based on a calibrated deep-learning model. However, regression-based models still require extensive ground-truth datasets for calibration, which are largely missing for most animal species.

The methods currently available in the scientific literature for egg size measurement still present limiting characteristics, such as manipulation of the eggs (calliper, water displacement, photogrammetry), ground-truthing (regression-based), and alignment with the subject (Silhouette). Moreover, most of these methods cannot be easily downscaled to microscopic conditions (light ranging or 3D point clouds).

This study directly addresses this knowledge gap by proposing a novel, non-contact optical method for the estimation of egg geometry and volume. Our methodology relies on commonly accessible materials (multiple-view angled photography and calibration markers) and an image processing routine, which we here refer to as the N-Views method. This low-cost and low-effort (computational and labour) method requires no physical contact with the egg, can be applied in field conditions and applies to avian and arthropod ovoids using digital cameras or microscopes. We present the hardware, computational methods and protocol required for such an estimation. We validate the N-Views method by comparing observations with manual methods (calliper and water buoyancy force), photogrammetry (similarly to Zhang et al., 2016), and the Silhouette method using samples from two avian species (*Gallus gallus domesticus* and *Coturnix coturnix*). We further describe the sensitivity of both the silhouette and the N-Views methods to two error sources (egg-misalignment and the effect of camera focal length), thus providing empirical evidence to support the design of optically-based egg size estimation protocols. Finally, we discuss the transferability of this protocol to the size estimation of microscopic arthropod eggs (the phytoseiid predatory mite *Amblyseius swirskii* Athias-Henriot). This research provides a robust, reproducible and open-source framework, overcoming limitations of previous methodologies.

## Materials and Methods

### Manual Measurements of egg geometries

Manual measurements were carried out on six *G. gallus domesticus* and six *C. coturnix* eggs. Breadth and Length were measured using an analog Vernier calliper with a resolution of 0.1 mm. Five independent measurements recording maximum length (L) and breadth (B) were conducted on each egg, and the mean value is reported. Egg volume was measured using the Archimedes’ buoyancy force method; for this, we measured the downward force exerted by each egg shape as introduced inside a distilled water beaker placed on a scale (0.001g resolution, LP620P Sartorius, Göttingen, Germany). The egg volume was estimated using the measured displaced water force and water density (0.001 g/cm3 accuracy DMA35 density meter, Anton Paar, Gratz, Austria).

### Silhouette method (monocular egg contour)

We used the Silhouette method (Soltani et al., 2015, Le Hesran et al., 2019) as the standard camera-based methodology to estimate geometries of *G. gallus domesticus, C. coturnix* and *A. swirskii* eggs. This method generally consists in retrieving a monocular top-view observation of the egg laid on a given surface (Figure 1A and Figure 2B). An estimation of the egg contour (silhouette) is produced by manual or automated segmentation from the background. In our case, we used a deep-learning segmentation method (described in section Deep-learning based egg segmentation routine). The major axis of the egg (Length) was estimated by applying Principal Component Analysis (PCA) to the contour coordinates and retrieving the first component direction. The maximum orthogonal radius to the main axis was also estimated, providing the egg maximum width (Breadth). The major axis divided the geometry in two roughly symmetrical sides (lobes). Using Pappus’ centroids theorem (assuming a solid of revolution symmetry), we estimated the Volume and Surface area as described in Le Hesran et al. (2019). Observations were taken from a nadir perspective, carefully placing the camera such that the sensor frame was as parallel to the egg main axis as possible. A calibrated scale reference was laid next to the egg to retrieve a pixel-to-meter scale correction factor.

**Figure 1.**
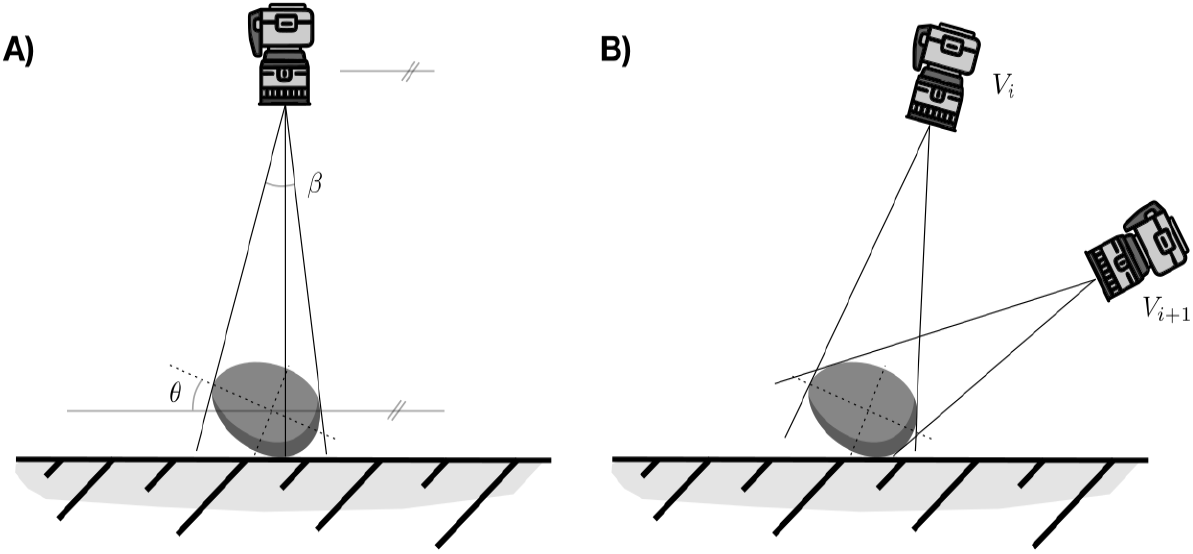
Scheme of non-orthogonal top views from an ovoid geometry (A) Silhouette method setup, (B) N-Views method setup.

**Figure 2.**
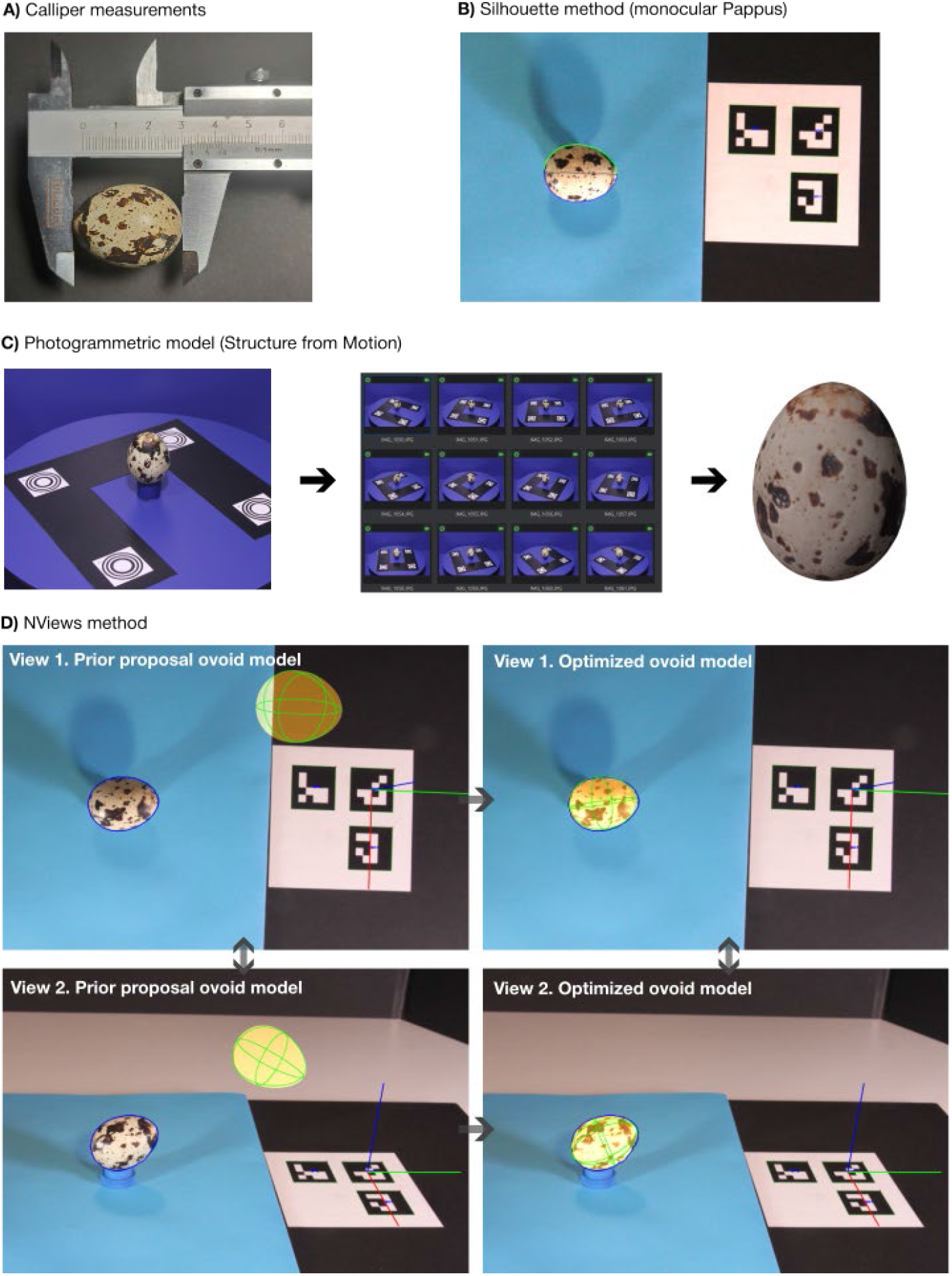
Example of four measurement methods for a *Coturnix coturnix* egg (CC-2) with A) Calliper, B) Silhouette method (green and blue contours depicting the detected right and left ovoid lobes), C) Photogrammetric reconstruction (structure from motion) and D) N-Views method ovoid fitting from two perspectives, depicting the real egg detection (SAM2) (blue contour), proposed/optimized virtual ovoid model mask (orange overlay), along with its main cross sections (green), and estimated camera-pose 3D coordinate system (red-greenblue axes) (left: egg model (prior) proposal, right: optimized egg model).

### N-Views ovoid estimation method

The novel proposed methodology to quantify geometrical characteristics of ovoid eggs (N-Views method) consists in acquiring a number (*N* ≈ [2, 12]) of arbitrarily oriented photographs from the targeted egg. A virtual (parametric) ovoid model is proposed in 3D, explicitly simulating the camera-lens characteristics and a perspective projection (see Figure 2D). The observed (real) egg contour is automatically retrieved for each image using a deep-learning semantic-segmentation model. An optimization routine fits the virtual ovoid model that projects to all views as closely as possible to the measured projected contours. This method, unlike the silhouette estimate, explicitly models the angle distortion of the egg (Figure 1A, angle *θ*) and a perspective projection model (Figure 1A, angle *β*), thus mitigating the major drawbacks of current optical egg estimation approaches. In the following sections we describe in detail all components of the N-Views method.

#### Camera model and pose estimation

A number of images (10-15) from a standard calibration board (Supp. Materials Figure A1.A) are used to derive the intrinsic camera matrix (focal length and sensor offset) and a lens distortion model (radial-tangential), defining a camera model. Additionally, a fiduciary triad marker (Supp. Materials Figure A1.B, ArUco ids 0,1,2) is placed in the scene and used to estimate each camera view *V*_*i*_ relative pose. Pose estimation is performed by automatically detecting all marker corners 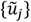, whose geometrical configurations are known (spacing and marker size). Corner coordinates are used to invert the world-to-camera pose, defined by each camera rotation (*r*_*i*_) and translation vectors (*t*_*i*_).

#### Deep-learning based egg segmentation routine

A binary mask 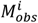 of the targeted egg at each camera view (*V*_*i*_) was extracted using a custom deep-learning segmentation tool built in Python (OpenCV4.9). This algorithm used the Segment Anything Model 2 (SAM2, Meta AI, *sam2_hiera_large*.*pt*) as a segmentation backbone (Ravi et al., 2024). The model is configured such that the user provides one or more points (few-shot inference mode) of the selected egg in view, and an automated segmentation is produced per image. The user can further refine the masks using positive or negative sample points and save the desired masks. All images were processed in a supervised manner, ensuring that segmentation adhered to the intended ovoid boundaries.

#### Parametric ovoid model

We adhere to the well-founded Preston Ovoid Equation (Preston, 1953), which has been reported to be a valid parameterization of natural egg ovoid shapes across multiple bird species. In particular, we adopt the Explicit Preston Equation (EPE) as presented by Shi et al., (2023). This is a five-parameter equation representing the contour of an egg around its local z-axis, defining a smooth, asymmetrical radial profile expressed as:

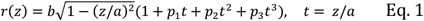

with *z* ∈ [−*a, a*], and parameters *p* = (*p*_1_, *p*_2_, *p*_3_), where *a* > 0 is the half-length, *b* > 0 is the equatorial half-breadth, and the parameters *p* modulate skewness and pointiness (asymmetry between blunt and sharp ends). The egg’s maximum length (*L* = 2*a*) and breadth (*B* = 2 ⋅ *max*_*z* ∈[−*a,a*]_*r*(*z*)) are then computed from the egg’s functional representation. Volume (*V*_*vol*_) and shell surface area (*S*) are derived by directly integrating the solid of revolution along the *z-axis*, following:

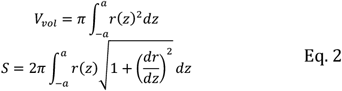

#### Ovoid model inversion

An automated optimization algorithm searches for a parametric egg model along its pose/orientation, such that the projection of such (virtual) egg matches as closely as possible all joint *N* camera observation views from the real ovoid. The egg model is here defined by 10 parameters, *x* = [*a, b, p*_1_, *p*_2_, *p*_3_, *t*_*x*_, *t*_*y*_, *t*_*z*_, *w*_*p*_, *w*_*y*_] ^*T*^, consisting in the five-parameter EPE egg profile (*a, b, p*_1_, *p*_2_, *p*_3_), a translation vector *t* = (*t*_*x*_, *t*_*y*_, *t*_*z*_) and a rotation scheme considering yaw (*w*_*y*_) and pitch (*w*_*p*_) rotations of the ovoid model. We did not include rotation around its own axis (roll), since it is unidentifiable given the proposed rotational symmetry ovoid model. The egg orientation vector was defined as:

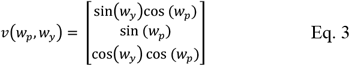

Equation 3 describes the world orientation of the local egg axis. In order to avoid gimbal lock singularities during the optimization process, we used a quaternion rotation *q*_*Δ*_(*a* → *v*) routine, representing the rotation of a local canonical orientation *a* = (0,0,1) to reach the target orientation vector *v*, with:

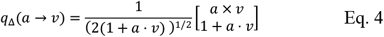

and with rotations being applied from an initial quaternion orientation *q*_0_ (e.g. egg local z-axis laying on the ground aligned with world axis-y) as: *q* = *q*_*Δ*_ ⊗ *q*_0_. The final egg rotation matrix *R*_*egg*_ was extracted from the unit quaternion as: *R*_*egg*_ = *RRt*(*q*).

The 3D surface of the proposed egg is then discretized on a grid in egg local cylindrical coordinates with azimuth *ϕ* and axial coordinate *z* whereas the meridional radius *r*(*z*) is given by the ovoid model (Eq. 1), thus producing a series of surface points on the virtual egg:

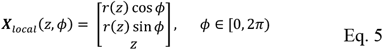

these surface points are transformed to world coordinates following a rotation-translation transform: *X*_*w*_ = *R*_*egg*_ *X*_*local*_ + *t*_*egg*_. Points from the proposed egg geometry in world coordinates, *X*_*w*_, are projected to each camera frame following the camera intrinsic, distortion and pose parameters and a pin-hole perspective camera model. We used the convex hull of these proposed projected ovoid points and rasterized them as a binary mask 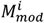, for each of the camera views (*V*_*i*_).

The difference between the real observations and the proposed egg shape and orientation is described by computing three complementary distance metrics between each egg model camera projection 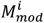 and the actually observed 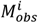:

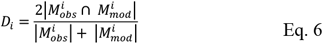

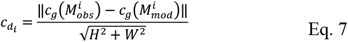

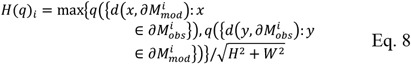

*D*_*i*_ is the DICE distance function, describing how closely both masks overlap, the distance 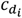 represents the distance between estimated and measured mask centroids (*c*_*e*_), and *H*(*q*)_*i*_ is the bidirectional quantile-Hausdorff distance between the edges of the observed and virtual masks, with *d*(*m, E*) the minimum of the distances between an arbitrary point x and all samples of the edge (E) and *q*(⋅) the quantile over a set. 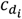 and *H*(*q*)_*i*_ are further normalized by the image diagonal 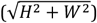. These terms are composed together in an objective function to be minimized:

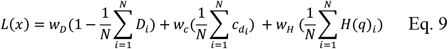

This loss represents the mean of all *N* views distance metrics in a weighted sum (*w*_*D*_ = 1, *w*_*C*_ = 1, *w*_*H*_ = 10), meanwhile the *H*(*q*) used the 90 percentile. The centroid distance serves to correct for poor (non-overlapping) egg proposals, the DICE distance allows the model to fine-tune to the observed masks once converging to a proximal estimate, meanwhile the Hausdorff distance is sensitive to the countour fit, helping the algorithm to adjust to the egg tip.

*L*(*m*) was minimized using the Powell optimization algorithm (scikit-learn), accepting initial and boundary conditions. The initialization values for the optimization vector 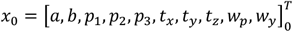 were derived using a simplified 3D model from all camera projections. Firstly, by estimating the translation using a multi-view triangulation of the centroid rays from all camera-mask views. Secondly, the major egg axis was estimated using a 2D PCA of the mask contours and from these, a least-squares problem was solved to find a common (initial) orientation parameter set [*w*_*p*_, *w*_*y*_]_0_ and main axes lengths [*a, b*]_0_, whereas the EPE asymmetry parameters were initialized as [*p*_1_, *p*_2_, *p*_3_]_0_ = 0.

### Photogrammetric 3D egg measurements

Macro-egg geometries were also measured with a photogrammetry approach (Figure 2C). This consisted in a 3D Structure from Motion (SfM) reconstruction setup in which ~200 images were taken per sample using a Canon EOS500D camera (Canon, Japan) and a 22 mm SIGMA 17-50mm 1:2.8 lens (Sigma, Japan). We used controlled lighting and background, fiduciary model scaling markers and a turntable. Some samples (e.g. smooth featureless eggs of *Gallus*) required painting visual features on their surface to allow for the SfM reconstruction. We used the software Meshroom (Griwodz et al., 2021) to perform a photogrammetric reconstruction routine and 3D model mesh estimation. This mesh was post-processed using MeshLab (eliminating surrounding non-egg geometries). This was further processed using a dedicated python script (based on trimesh and open3d libraries) to compute the longest possible axis length, width, volume and surface area from the 3D mesh.

#### *Gallus gallus domesticus* and *coturnix coturnix* egg geometry estimation and validation methods

A total of 12 unfertilized consumer-grade eggs were selected, six from *Gallus gallus domesticus* (GG) and six from *Coturnix coturnix* (CC). The inner content of each egg was replaced by a gypsum slurry, for preservation purposes. This was performed from a small access point created in the surface with a needle. The access point was smoothed out with gypsum, painted and sealed with a spray semi-mate varnish layer to avoid moisture exchange. Eggs were labelled as GG-1 to GG-6 and CC-1 to CC-6.

The geometrical characteristics of all eggs were derived using the four presented methods: Manual measurements Calliper/Archimedes (Length, Breadth and Volume), Photogrammetry (~200 images), N-Views (8 images) and Silhouette (1 image) (Length, Breadth, Volume and Surface area). During the measurements of N-Views, we used a lens (SIGMA 17-50 mm 1:2.8, Sigma, Japan) set to 50 mm focal length, mounted on a Canon EOS 500D (APS-C sensor, crop factor = 1.6, Canon, Japan), which corresponds to a 35 mm-equivalent focal length of 80 mm (we report hereinafter all focal lengths as 35 mm equivalent). We performed a best-effort Silhouette estimate using a relatively high focal length of 112 mm (70 mm set, CANON 55-250 mm 1:4-5.6, Canon, Japan) with approximate distance to targets of 90 cm from a top-view and carefully aligned egg main axis. Intrinsic calibration of the camera (focal length and distortion model) was done once per focal length setting. Additionally, two egg geometries (CC-2 and GG-2) were used to carry out three sensitivity tests:

**Test 1 - Number of views**: Evaluating the sensitivity of the N-Views method to varying numbers of acquired images (2, 4, 8 and 12 view observations with 10 replicates each). Samples were drawn randomly from a pool of 36 images (12 from near-nadir 80-90°, 12 from top-down with a 60° angle and 12 from a shallow 30° angle). Samples were distributed as follows: 2 views (1×90°, 1×60°), 4 views (2×90°, 1×60°, 1×30°), 8 views (3×90°, 3×60°, 2×30°), 12 views (4×90°, 4×60°, 4×30°).

**Test 2 - Specimen-axis misalignment**: Sensitivity of N-Views and Silhouette methods to varying distortions between the egg main axis and the camera sensor frame (Figure 1A, angle *θ*). Six deviations between 0° to 90° with the horizontal plane were tested.

**Test 3 - Focal length**: Sensitivity of N-Views and Silhouette methods to different focal length setups using a Canon EOS500D camera (crop factor 1.6, Canon, Japan) coupled with a focal length (35 mm equivalent) of 27 mm, 80 mm (17 mm and 50 mm set in a SIGMA 17-50mm 1:2.8 lens, Sigma, Japan) and 160 mm (100 mm set, CANON 55-250 mm 1:4-5.6, Canon, Japan).

#### Transference of the N-views method to the estimation of microscopic egg geometries

To showcase the application of the N-Views method to microscopic arthropod eggs, we used the phytoseiid mite *Ambylseius swirskii* (Acari: Phytoseiidae). This tiny arthropod (around 0.5 mm in length) is a generalist predator which is used worldwide for biological control of agricultural pests. *Amblyseius swirskii* females (Figure 3A) lay ovoid-shaped eggs (Figure 3B), which measure around 0.2 mm in length (Lopez, 2023). Two *A. swirskii* eggs were collected using a brush and laid on top of a calibration microscope slide. The markings in the microscope slide are used to estimate the pose of each view (acting as the fiduciary markers). Eight photographs of each sample were collected using a digital microscope (AD266S, Andonstar, Shenzhen, China) with an estimated focal length of 319 mm (35 mm equivalent sensor). Both the N-Views and the Silhouette measurement methods were used as described previously.

**Figure 3.**
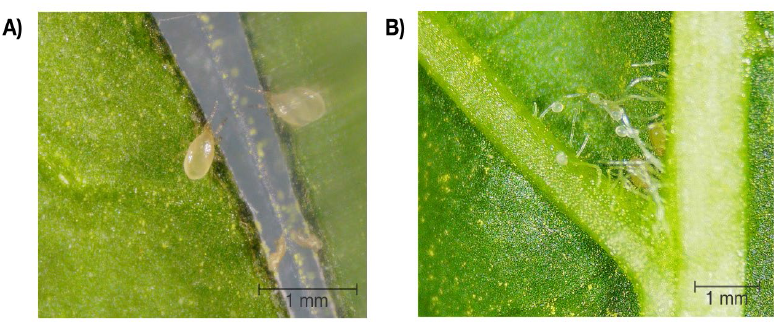
A) *Amblyseius swirskii* adult female, B) Acarodomatia on a sweet pepper leaf with a cluster of *A. swirskii* eggs.

## Results

Six *Coturnix coturnix* (CC-1 to CC-6) and six *Gallus gallus domesticus* (GG-1 to GG-6) eggs were measured using the four measurement methods described previously. A schema depicting the outputs of each method (A: Calliper, B: Silhouette (SIL), C: Photogrammetry (PG), and D: N-Views) for the sample egg CC-2 is presented in Figure 2.

Table 1 compares the results of the four methods across all samples for Length, Breadth and Volume (Appendix B in Suppl. Materials shows all comparisons and Normalized differences). We assume the Calliper method to be the standard reference for Length and Breadth estimates and the Archimedes buoyancy force measurements to be the reference for Volume estimations. Table 2 shows the mean absolute error between each method and its reference, along with the normalized relative error.

**Table 1.** Comparison of geometry measurements (Length, Breadth and Volume) with three optical methods (Photogrammetry, N-Views and Silhouette), 80 mm equivalent focal-length lens, compared with Calliper (Length and Breadth) and Archimedes (Volume) for 6 samples of *Coturnix coturnix* (CC) and 6 samples of *Gallus gallus domesticus* (GG) eggs.

| ID | Calliper |  | Arch<br>V<br>(mm<sup>3</sup>) | Photogrammetry |  |  | N-Views |  |  | Silhouette |  |  |
| --- | --- | --- | --- | --- | --- | --- | --- | --- | --- | --- | --- | --- |
|  | L<br>(mm) | B<br>(mm) |  | L<br>(mm) | B<br>(mm) | V<br>(mm<sup>3</sup>) | L<br>(mm) | B<br>(mm) | V<br>(mm<sup>3</sup>) | L<br>(mm) | B<br>(mm) | V<br>(mm<sup>3</sup>) |
| CC-1 | 32.80 | 25.22 | 10522.09 | 32.73 | 25.68 | 10507.48 | 32.43 | 25.18 | 10651.81 | 33.28 | 25.41 | 10994.89 |
| CC-2 | 33.02 | 24.99 | 10622.11 | 32.84 | 25.67 | 10459.22 | 32.90 | 25.06 | 10690.12 | 33.44 | 25.29 | 10971.02 |
| CC-3 | 33.02 | 25.11 | 10799.43 | 32.73 | 25.68 | 10507.48 | 32.55 | 25.10 | 10691.18 | 33.91 | 25.67 | 11277.08 |
| CC-4 | 33.76 | 24.48 | 10118.21 | 33.71 | 25.69 | 9973.61 | 33.70 | 24.33 | 10066.39 | 34.58 | 25.02 | 10638.66 |
| CC-5 | 34.00 | 24.06 | 9903.79 | 33.81 | 25.51 | 9682.19 | 33.33 | 23.99 | 9943.22 | 34.62 | 24.73 | 10444.39 |
| CC-6 | 33.00 | 26.04 | 11310.75 | 33.11 | 27.07 | 10507.48 | 33.07 | 25.98 | 11582.00 | 33.67 | 26.72 | 12100.52 |
| GG-1 | 53.62 | 39.86 | 44345.12 | 53.76 | 40.68 | 44458.56 | 53.40 | 39.66 | 44035.98 | 54.98 | 40.42 | 46538.04 |
| GG-2 | 50.00 | 39.98 | 41452.04 | 50.45 | 41.38 | 41204.03 | 50.01 | 39.93 | 41434.93 | 51.23 | 40.81 | 44073.94 |
| GG-3 | 51.06 | 41.06 | 44687.84 | 51.05 | 42.21 | 44248.63 | 50.97 | 41.02 | 44782.47 | 52.14 | 41.48 | 46130.84 |
| GG-4 | 54.86 | 45.14 | 57922.63 | 54.32 | 45.20 | 56279.47 | 54.83 | 45.00 | 57771.17 | 56.00 | 45.65 | 59652.09 |
| GG-5 | 48.26 | 39.44 | 38425.69 | 48.23 | 40.99 | 38217.08 | 48.27 | 39.12 | 38405.01 | 48.59 | 39.63 | 39502.32 |
| GG-6 | 57.10 | 44.76 | 59195.31 | 57.24 | 45.63 | 59996.11 | 57.26 | 44.71 | 59440.88 | 58.13 | 45.19 | 61369.35 |

**Table 2.** Summary statistics of geometry measurements errors (Length, Breadth and Volume) with three optical methods (Photogrammetry, N-Views and Silhouette). Mean Absolute Error and normalized mean absolute error (%) across 6 samples of *Coturnix coturnix* (CC) and 6 samples of *Gallus gallus domesticus* (GG) eggs. Length and Breadth are compared with Calliper measurements and Volume against Archimedes buoyancy force volume measurements.

| ID | Photogrammetry |  |  | N-Views |  |  | Silhouette |  |  |
| --- | --- | --- | --- | --- | --- | --- | --- | --- | --- |
|  | L<br>(mm) | B<br>(mm) | V<br>(mm<sup>3</sup>) | L<br>(mm) | B<br>(mm) | V<br>(mm<sup>3</sup>) | L<br>(mm) | B<br>(mm) | V<br>(mm<sup>3</sup>) |
| CC | 0.15<br>(0.4%) | 0.9<br>(3.6%) | 273.1<br>(2.5%) | 0.29<br>(0.88%) | 0.07<br>(0.27%) | 111.4<br>(1.03%) | 0.65<br>(1.95%) | 0.49<br>(1.96%) | 525.02<br>(4.96%) |
| GG | 0.22<br>(0.4%) | 0.97<br>(2.4%) | 575.5<br>(1.0%) | 0.09<br>(0.16%) | 0.13<br>(0.32%) | 139.7<br>(0.28%) | 1.03<br>(1.94%) | 0.49<br>(1.18%) | 1872.99<br>(3.99%) |

These results show a consistent submillimetre accuracy of the N-Views method across all measured samples (mean absolute difference with Calliper of 0.19 mm for Length and 0.10 mm for Breadth), whereas the Photogrammetry estimate showed a differential error profile (0.19 mm for Length and 0.94 mm for Breadth). Similarly, the Silhouette methodology overestimated all egg dimensions (0.84 mm for Length and 0.49 mm for Breadth). Across all tested methods, N-Views provided the best estimate of volume when compared with the reference measurements, resulting in a mean absolute error (MAE) of 125 mm^3^ (0.6%). Photogrammetry provided estimates with a mean absolute error of 424 mm^3^ (1.8%), and the Silhouette method of 1872 mm^3^ (4.4%).

During the measurement of all egg macro-samples (CC and GG), the N-Views method was tested using 8 camera observations of each egg. In order to test the sensitivity of this method to the number of camera orientations (views), we carried out a dedicated test (Test 1, described in Materials and Methods). Figure 4 shows the mean and two times the standard deviation (2*σ*) estimates of Length, Breadth, Volume and Surface of random 10 replicates for each of the four number of view sets. The reconstructions of Length and Breadth are robust to the number of views, with consistent deviations compared to Calliper observations and a dispersion 2*σ* (two times the standard deviation) for Length of [2 views: 0.32 mm, to 12 views: 0.18 mm], Breadth of [2 views: 0.29 mm, to 12 views: 0.28 mm], and Volume of [2 views: 336 mm^3^, to 12 views: 133 mm^3^].

**Figure 4.**
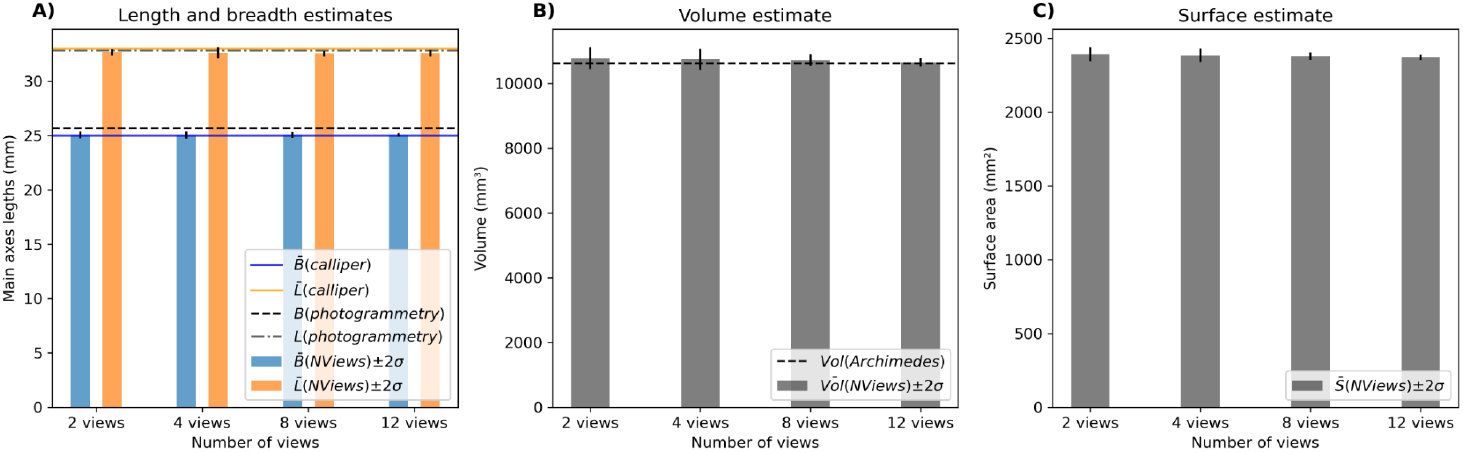
Effect of the number of images used during the N-Views reconstruction (mean and 2σ, across 10 replicates) on: A) Length and Breadth estimates, compared with those measured by Calliper and photogrammetry. B) Estimated volume and C) shell surface area (CC2 sample).

Figure 5 shows the sensitivity of geometrical estimates of the egg for variations in pitch (*θ*, see Figure 1A) angles ranging from 0° to 90° (Test 2). These results show that the N-Views method is largely independent (since sufficient information is captured from the N projections), whereas the Silhouette method is sensitive to the camera to egg main-axis alignment. Observations of Breadth through the Silhouette method remain largely constant (within the high observation error of ~ 2% for this method). On the other hand, estimations of Length changed rapidly with the observation angle. For instance, using the 0° value (of the Silhouette reconstruction) as reference, it showed deviations of [0.03, 0.08, 1.08, 2.78, 7.53] mm for CC-2 and [0.44, 0.51, 1.17, 2.9, 9.9] mm for GG-2 when having a distortion of [5.6°, 11.25°, 22.5°, 45° and 90° degrees] respectively. These results show that as the egg size increases, so does the sensitivity of the Silhouette method to axis-frame deviations, and that this error can be substantial (0.5 mm for even a relatively small deviation angle of 11.25°).

**Figure 5.**
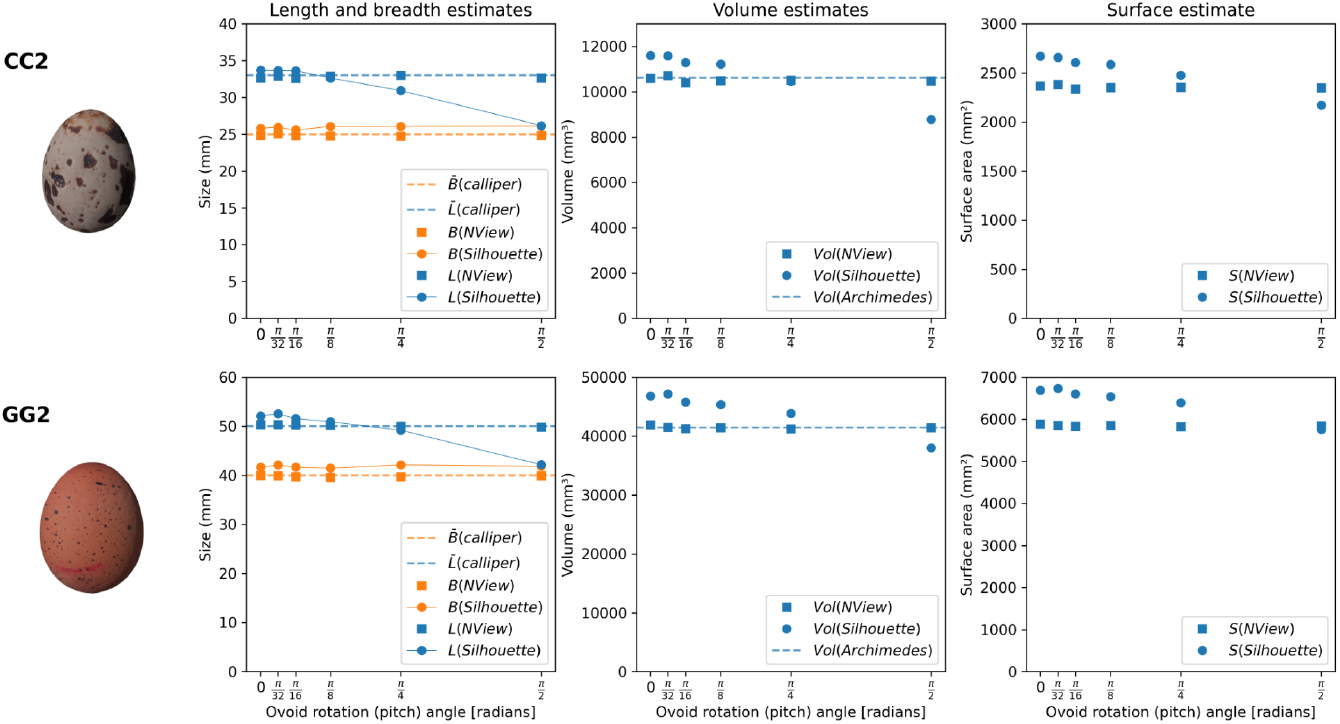
Effect of the ovoid pitch angle (0, 5.6, 11.25, 22.5, 45 and 90 degrees) with respect to the camera sensor for the N-Views method (squares) vs. Silhouette (circles). For comparison, dimensions measured by Calliper (Length and Breadth) and Archimedes (Volume) are provided. Above sample CC2, below sample GG2.

The camera projection angle *β* (Figure 1A), dominated by the effective focal length (*f*), also represents a source of error. In Table 3, we show the effect of increasing focal length in the Silhouette and N-Views methods (Test 3). While N-Views is relatively insensitive, the Silhouette method is shown to increase its accuracy as the focal length increases.

**Table 3.** Dependency of derived geometrical variables on focal length for the methods N-views and Silhouette (Sil) compared with the best estimates (Calliper and Archimedes) for two ovoid samples (CC2, GG2) aligned with the camera frame and under varying focal lengths (full frame equivalent) (27, 80, 160 mm), with an indicative distance camera-to-target.

| Id | Focal length (ff eq.) mm | Dist to target (cm) | Length (mm) |  |  | Breadth (mm) |  |  | Volume (mm<sup>3</sup>) |  |  | Surface area (mm<sup>2</sup>) |  |  |
| --- | --- | --- | --- | --- | --- | --- | --- | --- | --- | --- | --- | --- | --- | --- |
|  |  |  | Calliper | Nviews | Sil | Caliper | Nviews | Sil | Arch | Nviews | Sil | PG | Nviews | Sil |
| CC2 | 27 mm | 25 | 33.0 | 32.9 | 37.6 | 24.9 | 25.1 | 28.2 | 10622 | 10823 | 15287 | 2348 | 2399 | 3197 |
|  | 80 mm | 50 | 33.0 | 32.9 | 34.3 | 24.9 | 24.9 | 25.7 | 10622 | 10476 | 11491 | 2348 | 2347 | 2642 |
|  | 160 mm | 220 | 33.0 | 32.9 | 33.2 | 24.9 | 24.9 | 25.2 | 10622 | 10637 | 10658 | 2348 | 2372 | 2517 |
| GG2 | 27 mm | 25 | 50.0 | 51.2 | 56.9 | 39.9 | 40.0 | 45.6 | 41452 | 43202 | 60616 | 5837 | 6025 | 7960 |
|  | 80 mm | 50 | 50.0 | 50.0 | 52.9 | 39.9 | 39.7 | 42.4 | 41452 | 41188 | 49101 | 5837 | 5825 | 6918 |
|  | 160 mm | 220 | 50.0 | 50.1 | 50.8 | 39.9 | 39.8 | 40.6 | 41452 | 41519 | 42907 | 5837 | 5856 | 6334 |

We tested the transferability of the N-Views and Silhouette methods to the estimation of microscopic egg geometries using two samples: AS1 and AS2. Figure 6 shows two of the eight views used to reconstruct the sample AS2. Table 4 shows the comparison of estimated variables using the N-Views and Silhouette methods.

**Table 4.** Estimated parameters for two eggs of *Amblyseius swirskii*.

| ID | Method | L (mm) | D (mm) | V (mm<sup>3</sup>) | S (mm<sup>2</sup>) |
| --- | --- | --- | --- | --- | --- |
| AS-1 | N-Views | 0.2260 | 0.1614 | 0.003086 | 0.1044 |
|  | Silhouette | 0.2214 | 0.1660 | 0.003071 | 0.1111 |
| AS-2 | N-Views | 0.2196 | 0.1668 | 0.003190 | 0.1061 |
|  | Silhouette | 0.2206 | 0.1825 | 0.003306 | 0.1155 |

**Figure 6.**
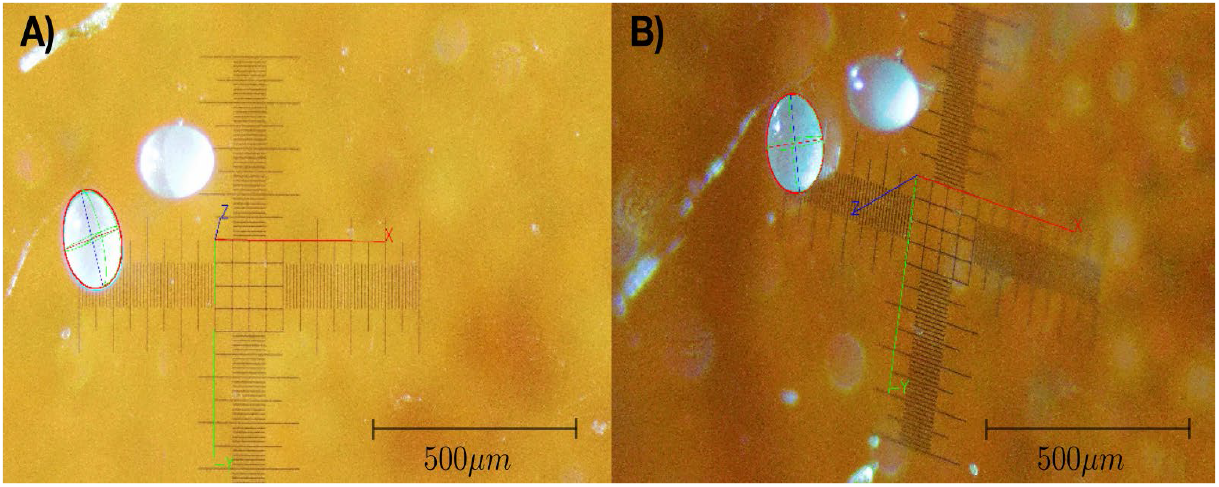
N-Views ovoid fit (red contour) estimate for *Amblyseius swirskii* egg sample (AS2) (2 views) along the estimated world-camera pose view (xyz axes).

## Discussion

The accurate and non-intrusive measurement of egg geometry represents a challenge in ecological research. This study introduced and validated the N-Views method, a novel computational approach that reconstructs a three-dimensional parametric model of an ovoid from a small number of arbitrarily oriented photographs. By explicitly modelling camera perspective and pose, our method overcomes the limitations of traditional monocular techniques. Across 12 samples (eggs from Coturnix and Gallus species), the N-Views method achieved sub-millimetre (~0.2 mm) agreement with calliper measurements in Length and Breadth, and relative errors of 1% and 0.28% for CC and GG in Volume measurements (mean absolute error of 125 mm^3^). This represents a quantitative accuracy improvement of 2-5 times compared to the results obtained with photogrammetry and monocular retrieval (Silhouette) in our tests.

Previous studies have presented different optical methods for egg volume quantification. For instance, for poultry eggs, Zhou et al. (2009) presented a Silhouette type strategy, reporting that 48% of their samples had an error larger than 1000 mm^3^. Similarly, Soltani et al. (2015) presented a Silhouette method (monocular approach with a calibrated neural network correction), reporting MAE = 590 mm^3^ (approx. 1% relative error), although their volumes were derived from an assumed egg density and not measured directly. Shi et al., (2023) proposed the EPE egg model and used the Silhouette method (EPE fit to a monocular egg view using a smartphone camera), where they reported a MAE of 1655 mm^3^ (1.5%) for *Anser cygnoides*, and MAE of 572 mm^3^ (2.2%) for *Phasianus colchicus* egg samples. Zhang et al. (2016) used a photogrammetry-based method reporting MAE of 584 mm^3^ in poultry eggs, whereas Okinda et al. (2020) reported an RMSE of 1175 mm^3^ and a 3.5% relative error when using a depth-camera approach to measure poultry eggs. Xiao et al. (2025) reported a MAE of 571 mm^3^ for duck eggs using the deep learning based regression method. In contrast, we report here mean absolute (and relative) error estimates for poultry eggs using the methods N-Views = 139.7 mm^3^ (0.28 %), Photogrammetry = 575 mm^3^ (1.0%) and Silhouette = 1873 mm^3^ (3.9%).

Our results show that the N-Views method is significantly more accurate than the Silhouette monocular approach and provides better results at a fraction of the operational burden required during Photogrammetric Structure from Motion (SfM) 3D reconstructions. Photogrammetry is a computationally intensive process, requiring a special setup, manipulation of the egg, the acquisition of 100-300 images from different angles, and the computation of a SfM (dense) reconstruction, which can take on the order of hours per sample. Moreover the photogrammetry method is only applicable if sufficient optical features from the surface of the egg can be retrieved (texture dependent).

The Silhouette method is more straightforward, requiring only one top-view image, the segmentation of a single image and a reference scale. However, the Silhouette method still results in substantial deviations compared to manual measurements. Two critical sources of error can explain these deviations: egg orientation (i.e., the egg’s major axis not being aligned with the camera’s optical plane, see Figure 1A, angle *θ*) and perspective distortion (particularly pronounced with the short focal length lenses often used in field photography, see Figure 1A, angle *β*). These effects have been quantified in Figure 5 and Table 3. The first error source could be tackled by very carefully orienting the egg, yet this is not trivial in practice.

The perspective error (angle *β*) arises from two possible sources. One is the fact that the optical ray intersects the ovoid at a given angle not representative of the maximum dimension (this is dependent on the curvature and size of the ovoid). Secondly, if the calibration marker in view is not coplanar with the egg mid-plane, this results in a magnification bias. For a conventional pin-hole camera, the lateral magnification relates as *m* ≈ *f*/*z*, with axial distance to the target *z* and the focal length *f*. A small Δ*z* change results in a magnification change of 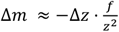, or in relative terms: 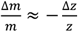, under constant framing of the egg in the image, *z* ∝ *f* and for a fixed Δ*z*, the magnification bias scales as ∝ 1/*f*. As a consequence, one would expect a reduction of this error source with increasing focal length, as seen in Table 3. However, when measuring macroscopic eggs, mitigating this error by increasing the focal length also requires increasing the distance to the target, which rapidly becomes impractical. For example, a 160 mm (equivalent) focal length requires a distance to the target of 2.2 m in a conventional DSLR camera. A direct way of mitigating both camera projection error sources when using the Silhouette approach would be the use of object-space telecentric lenses, which create a near orthographic view (*β* ≈ 0). In microscopy applications (see e.g. Le Hesran et al., 2019), this error source is mitigated, due to the use of telecentric tube lenses and when not (e.g., stereomicroscopes), due to the very small 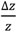 ratio. These error sources are directly tackled by the N-Views method by explicitly modelling the camera projection and the ovoid orientation from several views.

Furthermore, the proposed N-Views method allows for the recovery of egg geometries in a non-intrusive manner, since it does not require alignment of the camera and the object, and it requires no manipulation (e.g. calibration with calliper or submergence of the eggs). This is of interest for the practice of ecological studies in field conditions. Nevertheless, the possibility of retrieving egg geometries from real nests in the field is subject to having at least one view of optical access to the main axes of the egg. This may be achievable even if the eggs are laid in clusters, a common phenomenon in birds and arthropods (Faraji et al., 2002; Jetz et al., 2008), as long as the egg is laid horizontally within the nest (see example in Figure 8). Egg incubation posture in birds is influenced by many factors such as habitat, nest structure or egg shape (Birkhead et al., 2019; Hung et al., 2022). For example, the pyriform shape of some cliff-nesting birds’ eggs and their upright incubation posture are hypothesized to prevent eggs from rolling out of nests on cliff ledges (Hung et al., 2022). Depending on the bird species studied, the egg posture and position in the cluster might therefore affect our ability to measure them without contact. Regarding arthropod eggs, their posture depends on the species and their surrounding environment. In the case of *Amblyseius swirskii*, eggs are laid individually on non-glandular leaf trichomes or in clusters inside domatia, in a random posture (Lopez, 2023). Bird and arthropod eggs for which no visual access is available to the extremes of the egg main axis are still challenging to measure accurately. This limitation still prevails for all available optical methods if manipulation of the egg position is not possible. Further research on the sensitivity of the different methods to the retrieval of eggs laid in natural conditions should still be conducted.

**Figure 8.**
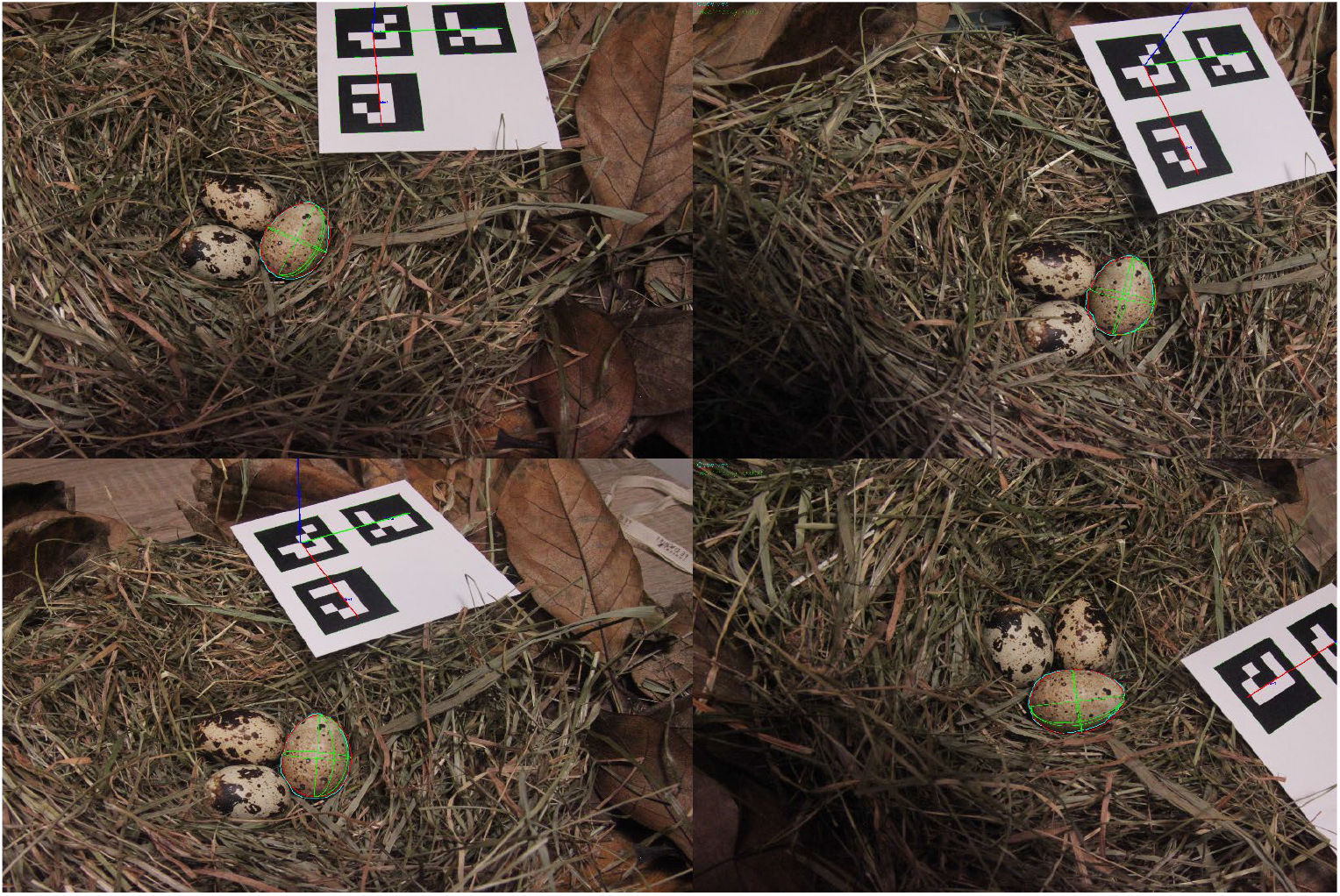
N-Views ovoid fit applied in a natural nest-like environment (example CC5).

The N-Views method still presents some limitations. For instance, it requires placing a scaling/pose retrieval (fiduciary) marker within the view of the captured photographs (see the example in Figure 8), which can be disruptive in natural environments. Furthermore, in the arthropod application displayed in this study, we used a standard calibration slide to perform pose estimations. The use of fiduciary markers (miniaturized) would facilitate and accelerate the processing. Additionally, when working at high magnifications, taking images at different angles while maintaining the markers and target in focus might be challenged by a reduced depth of field. In our example, this was however still achievable. Moreover, the tested SAM2 segmentation method to automate the retrieval of egg target masks proved relatively robust when testing natural-like environments (a bird nest). This should be further validated in a variety of environments with different egg characteristics. Alternatively, masks could be refined or directly manually delineated if needed. Another limitation of the N-Views method is the need to fit a parametric egg model. In this study, we used the Explicit Preston ovoid formulation from Shi et al., (2023), which has been reported as a flexible parametric functional egg shape that fits well with a broad range of avian species. However, this formulation does not account for egg deformations, non-revolution asymmetries, and non-ovoidal egg shapes and might therefore provide less accurate results in deformed or non-standard egg geometries.

## Conclusions

The N-Views approach produces a practical methodology for the measurement of egg dimensions, volume and surface area, without requiring direct contact with the ovoid (i.e. not requiring alignment), and no special hardware (just a consumer-grade camera and lens). Furthermore, this method is naturally extensible to microscopic eggs, as shown through its application to the measurement of *Amblyseius swirskii* eggs, and provides an effective protocol for biological studies independent on the scale (macro to microscopic applications). We found the N-Views method to be the most accurate source of egg volume determination with optical means (MAE = 125 mm^3^, 0.6% error) available in the scientific literature.

We further tested and evaluated the accuracy of other previously proposed methods for the optical determination of egg geometries (photogrammetry and monocular contour reconstructions/Silhouette), allowing to determine their relative accuracy and discuss the strengths and weaknesses of each approach. We demonstrated the dependency of the Silhouette method to different error sources (alignment and perspective), bringing new insights for the improvement of egg volume determination protocols and reporting.

Further research to test the retrieval capabilities of the N-Views method in natural conditions (i.e. nests for birds and leaves for arthropods) should still be conducted in the future, as this might be species and environment-dependent. Finally, we have quantified the accuracy of the different methods using Gallus and Coturnix eggs. The extension of the N-Views method to other species remains to be tested.

## Author contributions

Conceptualization: SLH, AMR; Methods implementation and software development: AMR; Investigation and analysis: AMR, SLH; Writing: SLH, AMR.

## Supplementary materials

## Appendix A. Calibration and Pose estimation markers

**Figure A1.**
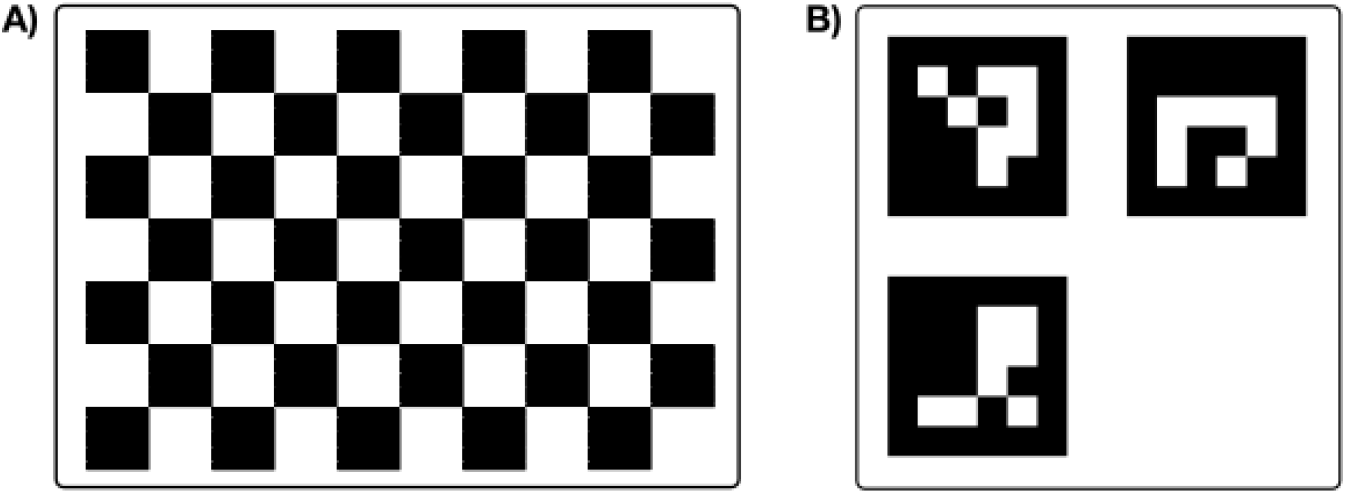
(A) Calibration board (estimation of camera intrinsics and distortion parameters), and (B) Triad of ArUco markers for camera extrinsic pose estimation.

## Appendix B. Additional Results

**Table S1.** Ovoid Length (L) estimates and error (normalized by Calliper measurements in %) for 6 samples of *Coturnix coturnix* (CC) and *Gallus gallus domesticus* (GG) using the methods Photogrammetry: PT, N-Views and Silhouette: Sil.

| ID | Length (mm) |  |  |  | Normalized difference (%) |  |  |
| --- | --- | --- | --- | --- | --- | --- | --- |
|  | Calliper | PT | N-Views | Sil | Calliper-PT | Calliper-N-Views | Calliper-Sil |
| CC1 | 32.80 | 32.73 | 32.43 | 33.28 | 0.21 | 1.13 | -1.45 |
| CC2 | 33.02 | 32.84 | 32.90 | 33.44 | 0.55 | 0.36 | -1.29 |
| CC3 | 33.02 | 32.73 | 32.55 | 33.91 | 0.88 | 1.43 | -2.69 |
| CC4 | 33.76 | 33.71 | 33.70 | 34.58 | 0.13 | 0.18 | -2.44 |
| CC5 | 34.00 | 33.81 | 33.33 | 34.62 | 0.55 | 1.96 | -1.82 |
| CC6 | 33.00 | 33.11 | 33.07 | 33.67 | -0.35 | -0.21 | -2.04 |
| GG1 | 53.62 | 53.76 | 53.40 | 54.98 | -0.25 | 0.42 | -2.53 |
| GG2 | 50.00 | 50.45 | 50.01 | 51.23 | -0.90 | -0.01 | -2.45 |
| GG3 | 51.06 | 51.05 | 50.97 | 52.14 | 0.01 | 0.17 | -2.12 |
| GG4 | 54.86 | 54.32 | 54.83 | 56.00 | 0.98 | 0.06 | -2.08 |
| GG5 | 48.26 | 48.23 | 48.27 | 48.59 | 0.06 | -0.02 | -0.68 |
| GG6 | 57.10 | 57.24 | 57.26 | 58.13 | -0.24 | -0.28 | -1.80 |

**Table S2.** Ovoid Breadth (B) estimates and error (normalized by Calliper measurements in %) for 6 samples of *Coturnix coturnix* (CC) and *Gallus gallus domesticus* (GG) using the methods Photogrammetry: PT, N-Views and Silhouette: Sil.

| ID | Breadth (mm) |  |  |  | Normalized difference (%) |  |  |
| --- | --- | --- | --- | --- | --- | --- | --- |
|  | Calliper | PT | N-Views | Sil | Calliper-PT | Calliper-N-Views | Calliper-Sil |
| CC1 | 25.22 | 25.68 | 25.18 | 25.41 | -1.84 | 0.17 | -0.74 |
| CC2 | 24.99 | 25.67 | 25.06 | 25.29 | -2.73 | -0.28 | -1.20 |
| CC3 | 25.11 | 25.68 | 25.10 | 25.67 | -2.29 | 0.05 | -2.25 |
| CC4 | 24.48 | 25.69 | 24.33 | 25.02 | -4.95 | 0.61 | -2.21 |
| CC5 | 24.06 | 25.51 | 23.99 | 24.73 | -6.02 | 0.31 | -2.79 |
| CC6 | 26.04 | 27.07 | 25.98 | 26.72 | -3.94 | 0.22 | -2.59 |
| GG1 | 39.86 | 40.68 | 39.66 | 40.42 | -2.04 | 0.50 | -1.40 |
| GG2 | 39.98 | 41.38 | 39.93 | 40.81 | -3.50 | 0.12 | -2.08 |
| GG3 | 41.06 | 42.21 | 41.02 | 41.48 | -2.79 | 0.10 | -1.03 |
| GG4 | 45.14 | 45.20 | 45.00 | 45.65 | -0.13 | 0.31 | -1.12 |
| GG5 | 39.44 | 40.99 | 39.12 | 39.63 | -3.94 | 0.81 | -0.48 |
| GG6 | 44.76 | 45.63 | 44.71 | 45.19 | -1.95 | 0.11 | -0.95 |

**Table S3.**
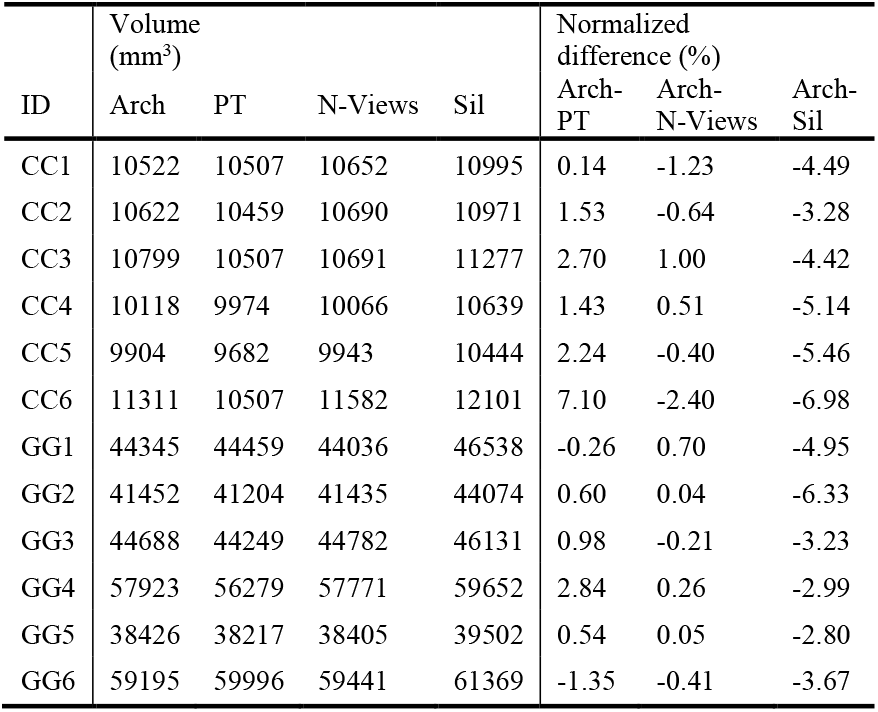
Ovoid Volume (Vol) estimates and error (normalized by Archimedes buoyancy measurements in %) for 6 samples of *Coturnix coturnix* (CC) and *Gallus gallus domesticus* (GG) using the methods Photogrammetry: PT, N-Views and Silhouette: Sil.

| ID | Volume (mm<sup>3</sup>) |  |  |  | Normalized difference (%) |  |  |
| --- | --- | --- | --- | --- | --- | --- | --- |
|  | Arch | PT | N-Views | Sil | Arch-PT | Arch-N-Views | Arch-Sil |
| CC1 | 10522 | 10507 | 10652 | 10995 | 0.14 | -1.23 | -4.49 |
| CC2 | 10622 | 10459 | 10690 | 10971 | 1.53 | -0.64 | -3.28 |
| CC3 | 10799 | 10507 | 10691 | 11277 | 2.70 | 1.00 | -4.42 |
| CC4 | 10118 | 9974 | 10066 | 10639 | 1.43 | 0.51 | -5.14 |
| CC5 | 9904 | 9682 | 9943 | 10444 | 2.24 | -0.40 | -5.46 |
| CC6 | 11311 | 10507 | 11582 | 12101 | 7.10 | -2.40 | -6.98 |
| GG1 | 44345 | 44459 | 44036 | 46538 | -0.26 | 0.70 | -4.95 |
| GG2 | 41452 | 41204 | 41435 | 44074 | 0.60 | 0.04 | -6.33 |
| GG3 | 44688 | 44249 | 44782 | 46131 | 0.98 | -0.21 | -3.23 |
| GG4 | 57923 | 56279 | 57771 | 59652 | 2.84 | 0.26 | -2.99 |
| GG5 | 38426 | 38217 | 38405 | 39502 | 0.54 | 0.05 | -2.80 |
| GG6 | 59195 | 59996 | 59441 | 61369 | -1.35 | -0.41 | -3.67 |

